# Targeting ACE2-RBD interaction as a platform for COVID19 therapeutics: Development and drug repurposing screen of an AlphaLISA proximity assay

**DOI:** 10.1101/2020.06.16.154708

**Authors:** Quinlin M. Hanson, Kelli M. Wilson, Min Shen, Zina Itkin, Richard T. Eastman, Paul Shinn, Matthew D. Hall

## Abstract

The COVID-19 pandemic, caused by SARS-CoV-2, is a pressing public health emergency garnering rapid response from scientists across the globe. Host cell invasion is initiated through direct binding of the viral spike protein to the host receptor angiotensin-converting enzyme 2 (ACE2). Disrupting the spike-ACE2 interaction is a potential therapeutic target for treating COVID-19. We have developed a proximity-based AlphaLISA assay to measure binding of SARS-CoV-2 spike protein Receptor Binding Domain (RBD) to ACE2. Utilizing this assay platform, a drug-repurposing screen against 3,384 small molecule drugs and pre-clinical compounds was performed, yielding 25 high-quality, small-molecule hits that can be evaluated in cell-based models. This established AlphaLISA RBD-ACE2 platform can facilitate evaluation of biologics or small molecules that can perturb this essential viral-host interaction to further the development of interventions to address the global health pandemic.

## Introduction

The SARS-CoV-2 pandemic has driven an urgent need to understand the molecular basis of infection and the consequent COVID-19 disease, and to rapidly identify pharmacologic interventions for both preventing infection and treating infected patients. Given that no approved therapeutics for treating any coronaviruses existed at the time SARS-CoV-2 emerged (late 2019), early attention has focused on drug repurposing opportunities ^1, 2^. Drug repurposing is an attractive approach to treating SARS-CoV-2, as active FDA-approved drugs or unapproved drug candidates previously shown to be safe in human clinical trials can be fast-tracked to the clinic. One such COVID-19 example is remdesivir (GS-5734, Gilead Sciences Inc.), a virus RNA-dependent RNA polymerase inhibitor that was in Phase II clinical trials for treating Ebola virus at the time SARS-CoV-2 emerged. Remdesivir was rapidly shown to be active against SARS-CoV-2 *in vitro*, and progressed to clinical trials leading to the FDA granting emergency use authorization ^3^. The understanding of each target and process critical for SARS-CoV-2 infection and replication will allow assays to be developed for both drug repurposing high-through screening, and to support new therapeutic development ^4^.

One therapeutic target receiving significant attention is the interaction between the SARS-CoV-2 spike protein (Uniprot – P0DTC2) and the host protein angiotensin-converting enzyme 2 (ACE2, Uniprot – Q9BYF1) ^5^. ACE2 is anchored to the extracellular surface of the cell and is responsible for catalyzing the conversion of angiotensin II into angiotensin 1-7, part of the renin-angiotensin system. The spike protein is exposed on the outer surface of the coronavirus particle, and has evolved to bind to ACE2 with high affinity ^6^. This binding between SARS-CoV-2 spike protein and ACE2 is the first step in viral infection ^7, 8^. The spike protein is comprises two functional regions: the S1 region contains the ACE2 receptor binding domain (RBD), and S2 region being responsible for membrane fusion. Because ACE2 binding by RBD is the first step in virus infection, preventing the interaction between Spike RBD and ACE2 is considered a viable therapeutic strategy, and work with prior zoonotic coronaviruses SARS and MERS has demonstrated proof-of concept for this approach ^9–11^. The primary therapeutic strategy being pursued for inhibiting the spike-ACE2 interaction is the development of selective antibodies against the spike RBD. Other approaches include antibodies that bind to ACE2 (though these may produce side-effects) ^12^, development of soluble recombinant ACE2 to compete for spike binding and limit cell entry ^13^, or small molecules that directly bind to ACE2 and biophysically reduce its affinity for binding to spike protein.

At NCATS, we are developing both protein/biochemical and cell-based assays to interrogate a number of biological targets to enable identification of potential therapeutic leads. Our initial focus is on performing drug repurposing screening for each assay and rapidly sharing the data through the NCATS OpenData portal for COVID-19 (https://opendata.ncats.nih.gov/covid19) ^14^. As part of this effort, we sought to develop a proximity assay for measuring the interaction between ACE2 and SARS-CoV-2 spike RBD to facilitate the identification of small molecules and biologics that disrupt this essential process.

AlphaLISA is a proximity-based assay that uses a pair of donor and acceptor beads to measure interaction (or disruption) of two tagged proteins/targets of interest ^15^. When the donor bead is excited by light at 680 nm it converts ambient oxygen to singlet oxygen. If an acceptor bead is in close proximity to the singlet oxygen, it emits a luminescent signal at 615 nm (and when this proximity is perturbed, there is a loss of signal). As AlphaLISA is amenable to quantitative high-throughput sceeening (qHTS), and can leverage multiple tagged ACE2 and Spike RBD constructs that are commercially available, this methodology could prove useful as a discovery approach for small or large molecules that inhibit infection ^15, 16^.

Herein we describe the development of an AlphaLISA assay using a recombinant SARS-CoV-2 spike protein RBD fused to a Fc tag and recombinant soluble human ACE2 fused to a biotinylated AviTag to model a simplified spike RBD-ACE2 interaction system. The ratio of proteins in the assay was optimized, and proof-of-concept protein-protein disruption demonstrated using untagged ACE2 or RBD protein as competitive inhibitors. The assay was used for a drug repurposing screen of >3K compounds, using a library of approved small molecule drugs (the NCATS Pharmaceutical Collection, NPC) and a library of compounds with prior demonstrated anti-infective activity ^17, 18^. We also screened all compounds in parallel against the TruHits counter-assay to identify and rule out false-positive hits.

## Methods

### Reagents

ACE2-His-Avi (human ACE2 residues 18-740, C-terminal His-Avi tags) (Cat # AC2H82E6) was acquired from ACROBiosystems (Newark, DE). RBD-Fc (SARS-CoV-2 spike protein residues 319-541, C-terminal Fc tag) (Cat #40592-V02H), ACE2-His (ACE2 residues 1-740, C-terminal His tag) (Cat # 10108-H08H), and S1-His (spike protein residues 1-667, C-terminal His tag) (Cat # 40150-V08B1) were acquired from Sino Biological (Wayne, PA). All proteins were reconstituted in 1X PBS at pH 7.4 supplemented with 0.05 mg/mL bovine serum albumin (BSA). Streptavidin Donor beads (Cat # 6760002) and Protein A acceptor beads (Cat # AL101) were acquired from PerkinElmer (Waltham, MA).

### Cross-titration experiment

Concentrations of ACE2-His-Avi and RBD-Fc in the AlphaLISA assay were selected by cross-titrating ACE2-His-Avi (300 - 0.1 nM) against RBD-Fc (300 – 0.1 nM) in 1X PBS (pH 7.4) supplemented with 0.05 mg/mL BSA. Each concentration combination of ACE2-His-Avi and RBD-Fc were mixed in a 384-well plate (square-well, high-base, white, medium binding; Cat # EWB010000A, Aurora Microplates, Whitefish, MT) and incubated at 25°C for 30 minutes. Streptavidin Donor beads and Protein A Acceptor beads were then added to the wells using a multi-channel pipet to a final concentration of 10 μg/mL each. AlphaLISA signal was read using a PheraSTAR (BMG Labtech, Cary, NC) plate reader with a 384-well format focal lens equipped with an AlphaLISA optical module (BMG Labtech, Cary, NC). The optimal concentration combination of ACE2-His-Avi and RBD-Fc was identified by the combination that returned the highest signal in the presence of 10 µg/mL Streptavidin Donor beads and 10 μg/mL Protein A Acceptor beads.

### Competition ACE2-and S1-binding assay

A competition assay was performed to confirm the AlphaLISA assay was sensitive the ACE2-His-Avi interactions with RBD-Fc. ACE2-His (Sino Biological, Wayne, PA) and S1-His (Sino Biological, Wayne, PA) are not recognized by either the Streptavidin Donor bead nor the Protein A Acceptor bead. A gradient of ACE2-His concentrations (1 µM to 0.01 nM) were mixed with RBD-Fc (4 nM). After 30 minutes of preincubation at 25 °C ACE2-His-Avi (4 nM) was added to the mixture and allowed to incubate at 25 °C for another 30 minutes. A mixture of Streptavidin donor beads and Protein A acceptor beads were then added to a final concentration of 10 µg/mL each. The resulting mixture was incubated at 25 °C for 40 minutes before reading the AlphaLISA luminescent signal with a PheraSTAR plate reader. S1-His competition assay was performed in the same manner with some changes S1-His (300 nM to 0.1 nM) was mixed with ACE2-His-Avi (4 nM) and preincubated for 30 minutes at 25 °C, then RBD-Fc was introduced to the mixture before adding beads.

### AlphaLISA Screening Protocol

The AlphaLISA screening assay was performed according to the protocol in Table 1.

**Table 1.** Detailed SARS-CoV-2 spike-ACE2 AlphaLISA qHTS protocol.

| Step # | Process | Notes |
| --- | --- | --- |
| 1 | 20 nL of compounds and controls were pre-dispensed into 1536-well plates. | Compound transfer performed using an ECHO 650 acoustic dispenser (LabCyte). 20 nL of compounds in dose response were transferred to columns 5-48. Positive control and DMSO (neutral control) were dispensed into Columns 1-4. 1536-well, square well, high base, white plate, solid bottom, non-sterile, non-treated; Cat # EWB010000A, Aurora Microplates. |
| 2 | Dispense 2 μL ACE2-His-Avi; to wells except column 2 | 4X ACE2-His-Avi (16 nM, Cat # AC2-H82E6, Acro Biosystems) in PBS (Cat # 10010-031, ThermoFisher) + 0.05 mg/mL BSA (Cat # BP1600-1, Fisher Scientific) |
| 3 | Incubate at RT for 30 minutes, 200 rpm shaking | Preincubation of ACE2 with compound step |
| 4 | Dispense 2 μL RBD-Fc to wells, except column 3 | 4X RBD-Fc (16 nM, Cat # 40592-V02H, Sino Biological) in PBS + 0.05 mg/mL BSA |
| 5 | Incubate at RT for 30 minutes, 200 rpm shaking | Equilibration of RBD-Fc and ACE2-Avi complexes |
| 6 | Dispense 4 μL of AlphaLISA Donor/Acceptor beads, except column 1 | 2X Protein A acceptor beads (20 μg/mL, Cat # AL101M, PerkinElmer), 2X Strep donor beads (20 μg/mL, Cat # 6760002, PerkinElmer) in PBS + 0.05 mg/mL BSA |
| 7 | Incubate at RT for 30 minutes, 200 rpm shaking | Final assay conditions are 4 nM ACE2-His-Avi, 4 nM S1-RBD-Fc, 1X PBS, 0.05 mg/mL BSA, 10 μg/mL Protein A Acceptor beads, 10 μg/mL Streptavidin donor beads |
| 8 | Read on PHERAstar FSX (BMG Labtech) | Fastest read settings, AlphaLISA Module (680 nm excitation, 615 nm emission) (Cat # 1802H1, BMG Labtech) |

### TruHits Counter-assay

The TruHits counter-assay was performed according to the protocol Table 2.

**Table 2.** Detailed TruHits AlphaLISA counter-assay protocol.

| Step # | Process | Notes |
| --- | --- | --- |
| 1 | 20 nL of compounds and controls were pre-dispensed into 1536-well plates. | Compound transfer performed using an ECHO 650 acoustic dispenser (LabCyte); compounds in dose to columns 5-48, controls to columns 1-1536-well plates, Cat # EWB010000A, Aurora Microplates. |
| 2 | Dispense 3 μL Biotin-BSA Acceptor Beads to all wells, except column 2. | 2X Biotin-BSA Acceptor beads (0.02 mg/mL, TruHits kit Cat # 6760627, PerkinElmer) in PBS (Cat # 10010-031, ThermoFisher) + 0.05 mg/mL BSA (Cat # BP1600-1, Fisher Scientific) |
| 3 | Incubate at RT for 30 minutes, 200 rpm shaking | Incubation of Biotin-BSA acceptor beads with test compounds. |
| 4 | Dispense 3 μL Streptavidin Donor beads to wells, except column 3 | 2X Streptavidin Donor beads (0.02 mg/mL, TruHits kit Cat # 6760627, PerkinElmer) in PBS (Cat # 10010-031, ThermoFisher) in PBS + 0.05 mg/mL BSA |
| 5 | Incubate at RT for 30 minutes, 200 rpm shaking | Streptavidin-Biotin complex formation. Final assay conditions: 10 μg/mL Streptavidin donor beads, 10 μg/mL Biotin-BSA acceptor beads in PBS + 0.05 mg/mL BSA. |
| 6 | Read on PHERAstar FSX (BMG Labtech) | Fastest read settings, AlphaLISA Module (680 nm excitation, 615 nm emission) (Cat # 1802H1, BMG Labtech) |

### Data Processing and Analysis

To determine compound activity in the qHTS assay, the concentration-response data for each sample was plotted and modeled by a four-parameter logistic fit yielding IC_50_ and efficacy (maximal response) values. Raw plate reads for each titration point were first normalized relative to positive control (-100% activity, full inhibition) and DMSO-only wells (basal, 0% activity). Data normalization and curve fitting were performed using in-house informatics tools. Compounds were designated as Class 1–4 according to the type of concentration–response curve (CRC) observed ^19^. In brief, Class −1.1 and −1.2 were the highest-confidence complete CRCs containing upper and lower asymptotes with efficacies ≥ 80% and < 80%, respectively. Class −2.1 and −2.2 were incomplete CRCs having only one asymptote with efficacy ≥ 80% and < 80%, respectively. Class −3 CRCs showed activity at only the highest concentration or were poorly fit. Class 4 CRCs were inactive having a curve-fit of insufficient efficacy or lacking a fit altogether. Data visualization was performed using Prism (GraphPad, San Diego, CA) and Spotfire (PerkinElmer) software programs. Scheme figures were all prepared using Illustrator (Adobe, San Jose, CA).

## Results

### Assay Design

We set out to develop a 1536-well plate assay capable of measuring SARS-CoV-2 spike protein receptor binding domain (RBD) to the ACE2 receptor. Two recombinant protein constructs were selected to model the RBD-ACE2 interaction *in vitro*. A truncated spike protein RBD (residues 319-541) fused to a C-terminal Fc tag and a truncated soluble ACE2 (residues 18-740) fused to a C-terminal poly-His tag followed by an AviTag (Figure 1A). These constructs were selected because the Fc tag and AviTag form tight and specific interactions with Protein A and streptavidin, respectively, making them ideal handles for AlphaLISA particles. When the RBD binds to ACE2 the Fc-tag and AviTag are accessible to Protein A acceptor beads and Streptavidin donor beads, respectively (Figure 1B). This series of interaction brings the donor and acceptor beads into sufficient proximity to excite the donor bead at 680 nm and monitor acceptor bead luminescence at 615 nm.

**Figure 1:**
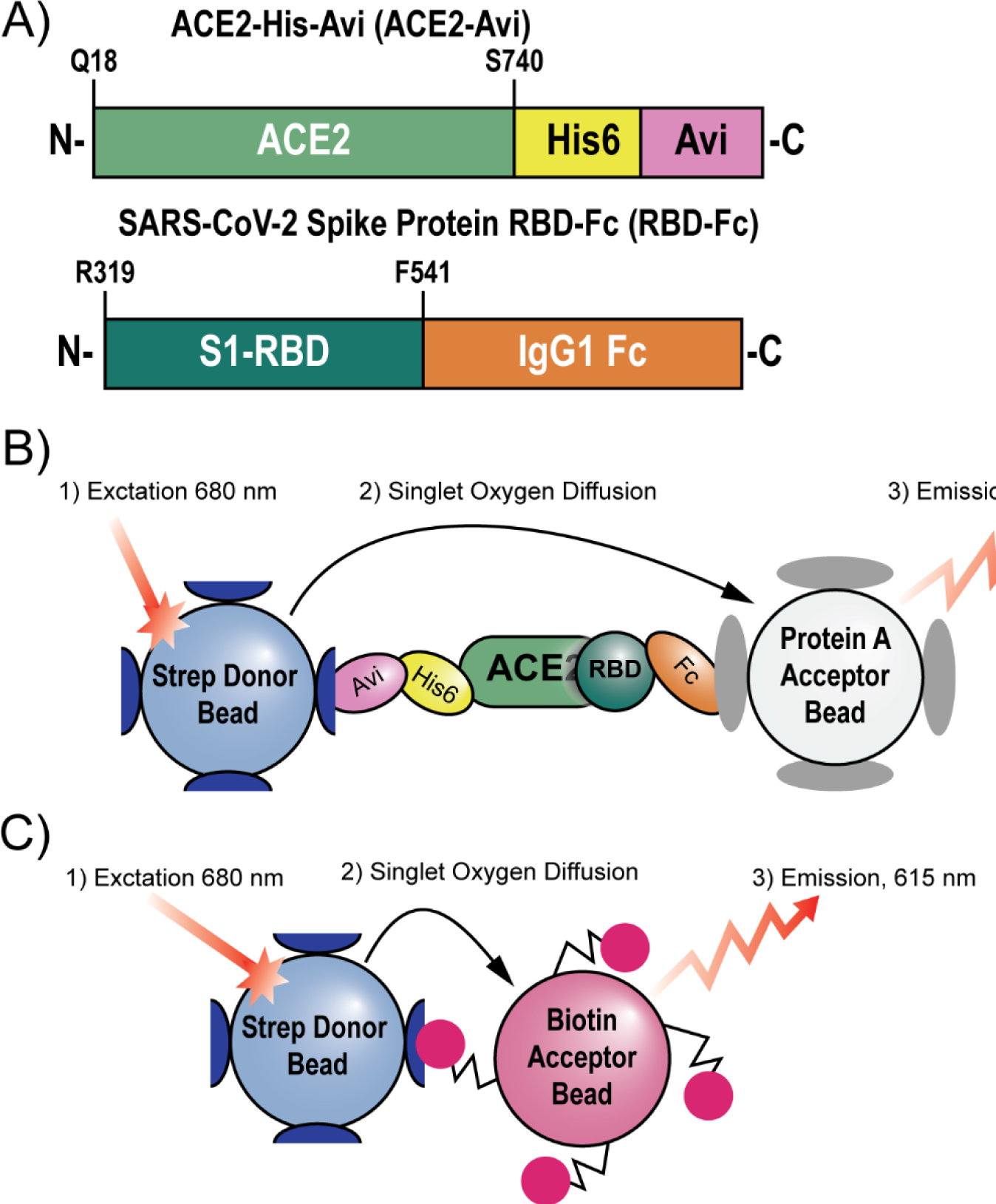
Scheme describing the assays employed in this paper. A) The recombinant protein constructs ACE2-His-Avi (ACE2-Avi) (Acro Biosystems) and SARS-CoV-2 Spike Protein Receptor Binding Domain-Fc (RBD-Fc) (Sino Biological) were used to model ACE2-RBD binding. B) AlphaLISA assay system used to monitor ACE2-RBD interacts. Streptavidin donor beads recognize the Avi tag on ACE2. Protein A acceptor beads recognize the Fc tag on RBD. When in proximity the donor beads can be excited with light at 680 nm. This generates singlet oxygen which diffuses to the acceptor beads, causing the acceptor beads to luminesce at 615 nm. C) The TruHits counter-screen uses strep donor beads which directly interact with biotin acceptor beads. Because no intermediary molecule is needed to bring the donor and acceptor beads in proximity, the TruHits assay can identify compounds which directly interfere with the AlphaLISA readout.

### Defining Assay Parameters

We first had to determine the working concentrations of RBD-Fc and ACE2-Avi that maximize the AlphaLISA signal. 10 µg/mL was selected for the concentration of donor and acceptor beads because it gave good signal with minimal well-to-well cross-talk (data not shown). To determine the optimal protein concentrations for the assay, we performed a cross-titration of RBD-Fc and ACE2-Avi varying each from 300 nM to 0.1 nM (Figure 2A). The maximum signal was achieved at 3.7 nM of both RBD-Fc and ACE2-Avi. Next, we tested the signal consistency of a high-signal control (ACE2-Avi + RBD-Fc + both AlphaLISA beads) and a low-signal control (ACE2-Avi + RBD-Fc + streptavidin donor beads) in 1536-well format (Figure 2B). At 4 nM of ACE2-Avi, 4 nM RBD-Fc, and 10 µg/mL of both streptavidin donor beads and Protein A acceptor beads we determined a plate Z’ value of 0.73 and signal-to-background of 286.7, indicating satisfactory assay performance.

**Figure 2:**
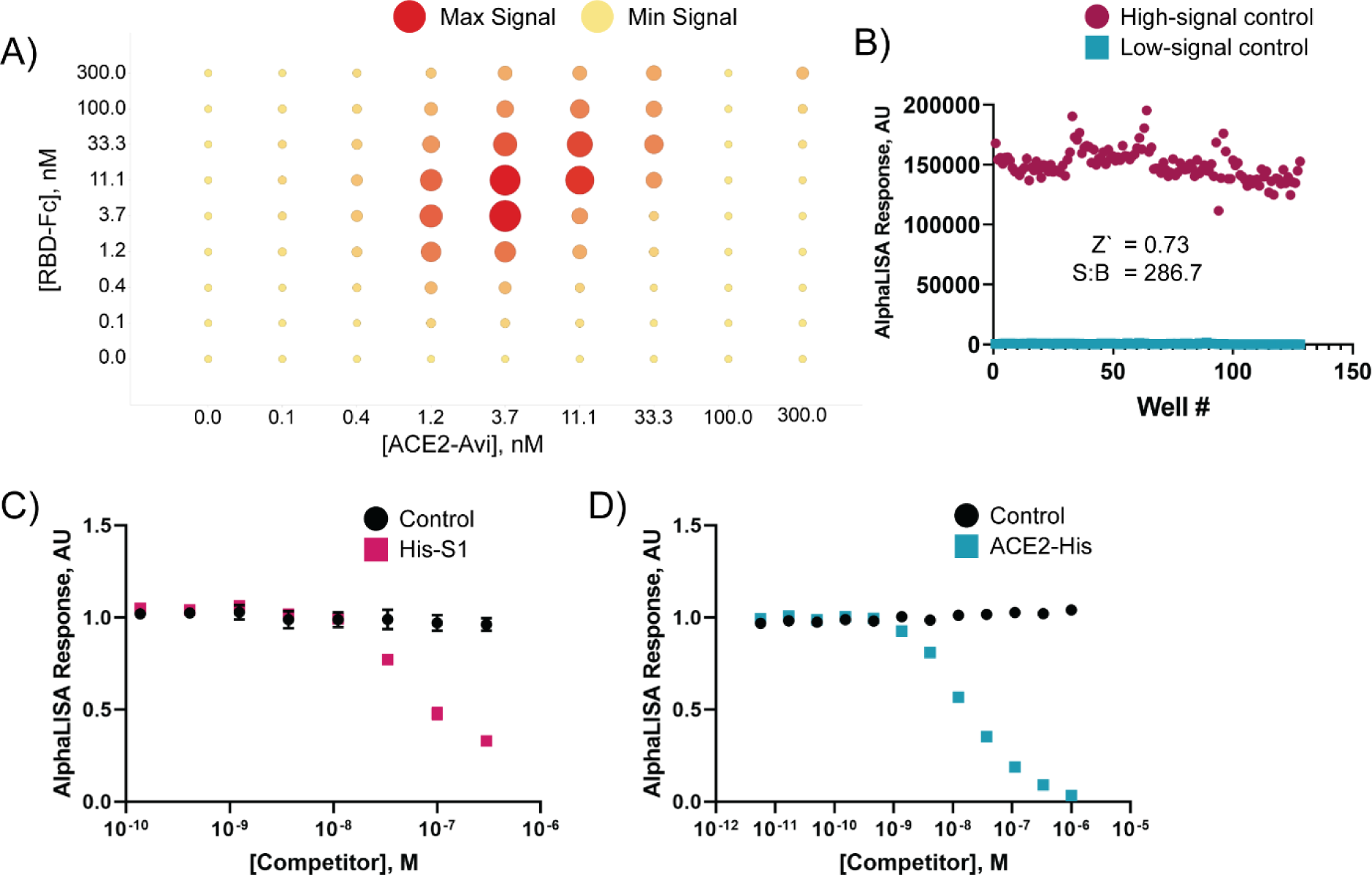
The AlphaLISA assay monitors ACE2-RBD interactions. A) ACE2-Avi and RBD-Fc were titrated against each other from 300 – 0.1 nM in matrix format to determine the optimal protein concentrations for the AlphaLISA assay. ACE2-Avi and RBD-Fc were mixed at the concentrations indicated and allowed to equilibrate at 25°C for 30 minutes. Streptavidin Donor beads and Protein A Acceptor beads were then introduced to the solutions to a final concentration of 5 µg/mL of each bead. After 40 minutes of incubation at 25°C the signal intensity was read using a PheraStar plate reader. B) Assay suitability to 1536-well format was determined by combining 4 nM ACE2-Avi with 4 nM RBD-Fc in PBS + 0.05 mg/mL BSA. The mixture was incubated at 25 °C for 30 minutes. Streptavidin donor beads (10 µg/mL) were added to columns 1-8 of a 1536-well plate. Protein A acceptor beads (10 µg/mL) were added to columns 5-8. The entire mixture was incubated at 25°C for 40 minutes before reading the low-signal control (columns 1-4) and high-signal control (columns 5-8) using a PheraSTAR plate reader with an AlphaLISA module. Z’ and signal-to-background (S:B) were calculated from these sample measurements. C, D) To confirm the AlphaLISA signal related to ACE2-RBD interactions, we tested the ability of His-S1 (another form of the SARS-CoV-2 Spike protein) and untagged ACE2 to lower the AlphaLISA signal in a dose-dependent manner. C) His-S1 was mixed in dose-response with 4 nM ACE2-Avi and allowed to incubate at 25°C for 30 minutes before adding 4 nM RBD-Fc. D) Untagged ACE2 was pre-incubated in dose-response with 4 nM RBD-Fc and allowed to incubate at 25°C for 30 minutes before adding 4 nM ACE2-Avi. Both His-S1 and untagged ACE2 showed dose-dependent signal loss, indicating the AlphaLISA signal is mediated by RBD-Fc binding to ACE2-Avi.

Next we wanted to confirm the AlphaLISA signal measured was dependent on the RBD-Fc interaction with ACE2-Avi (i.e., the assay is ‘inhibitable’). We tested the ability of a truncated SARS-CoV-2 spike protein with a C-terminal poly-His tag to compete with RBD-Fc for ACE2 binding, and a soluble ACE2 with a C-terminal poly-His tag to compete with ACE2-Avi for RBD binding. These constructs were selected because both the S1-His and ACE2-His constructs should recognize their respective binding partner, ACE2-Avi and RBD-Fc, but will not be engaged by the AlphaLISA beads. Indeed, both S1-His and ACE2 were able to compete with their respective binding partners in a dose-dependent manner (Figure 2C, D), indicating that the AlphaLISA assay is dependent on RBD-ACE2 interaction, and therefore ‘inhibitable’.

### qHTS screen of NPC and AIL compounds sets

Having established the conditions and suitable performance for the RBD-ACE2 AlphaLISA assay, we employed it for qHTS. To this end we selected two libraries: the NCATS Pharmaceutical Collection (NPC, small-molecule library of approved drugs in the USA, Europe and Japan, 2,678 compounds), and an in-house anti-infectives library (AIL, compounds shown to be active against infectious diseases, 739 compounds) to screen against our AlphaLISA RBD-ACE2 binding assay (HTS assay protocol shown in Table 1). When screening the NPC library the average plate Z’ was 0.77 with an average signal-to-background of 108.5 (Figure 3A). The AIL collection screen had an average plate Z’ of 0.90 and signal-to-background of 460.7 (Figure 3B). Both NPC and AIL were also screened in parallel against the TruHits counter-assay platform to rule out false-positive hits (Figure 1C, TruHits HTS protocol displayed in Table 2). For NPC, the TruHits counter-assay performed with an average Z’ of 0.93 and average signal-to-background of 15477 (Figure 3C) and the AIL had an average Z’ of 0.94 with an average signal-to-background of 17,410 (Figure 3D).

**Figure 3:**
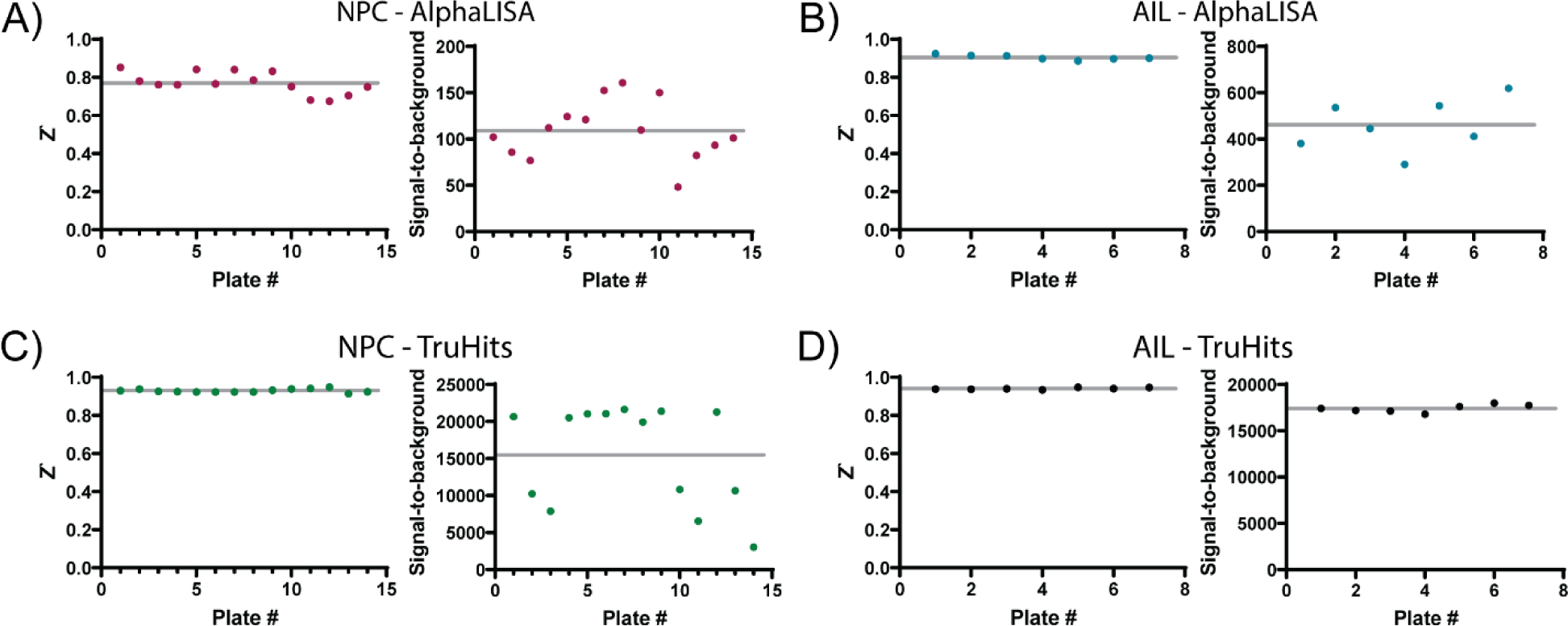
Assay performance for compound libraries tested in the primary AlphaLISA assay and the TruHits counter-assay. Z’ scores and signal-to-background values are plotted for each plate in the NPC AlphaLISA assay (A), the AIL AlphaLISA Assay (B), the NPC TruHits counter-screen (C), and the AIL TruHits counter-screen (D). Grey lines in all plots represent the mean values.

### Hit Identification

Hit compounds for both the AlphaLISA primary assay and the TruHits counter screen were identified as those with an IC_50_ < 50 µM and a maximum efficacy ≥ −50%, effectively removing all class −4 and some class −3 compounds (see Methods for curve class designations). A total of 574 initial hits were identified for the AlphaLISA assay (16.9% hit rate) and a total of 624 initial hits for the TruHits counter-assay (18.4% hit rate, Figure 4). 89 of these initial hits were unique to the AlphaLISA assay forming our first pool of potential active compounds. High-quality hits were then selected by visual inspection of the dose-response curves (Figure 5). 25 compounds were identified as high-quality hits capable of disrupting RBD-ACE2 binding (Table 3).

**Table 3.**
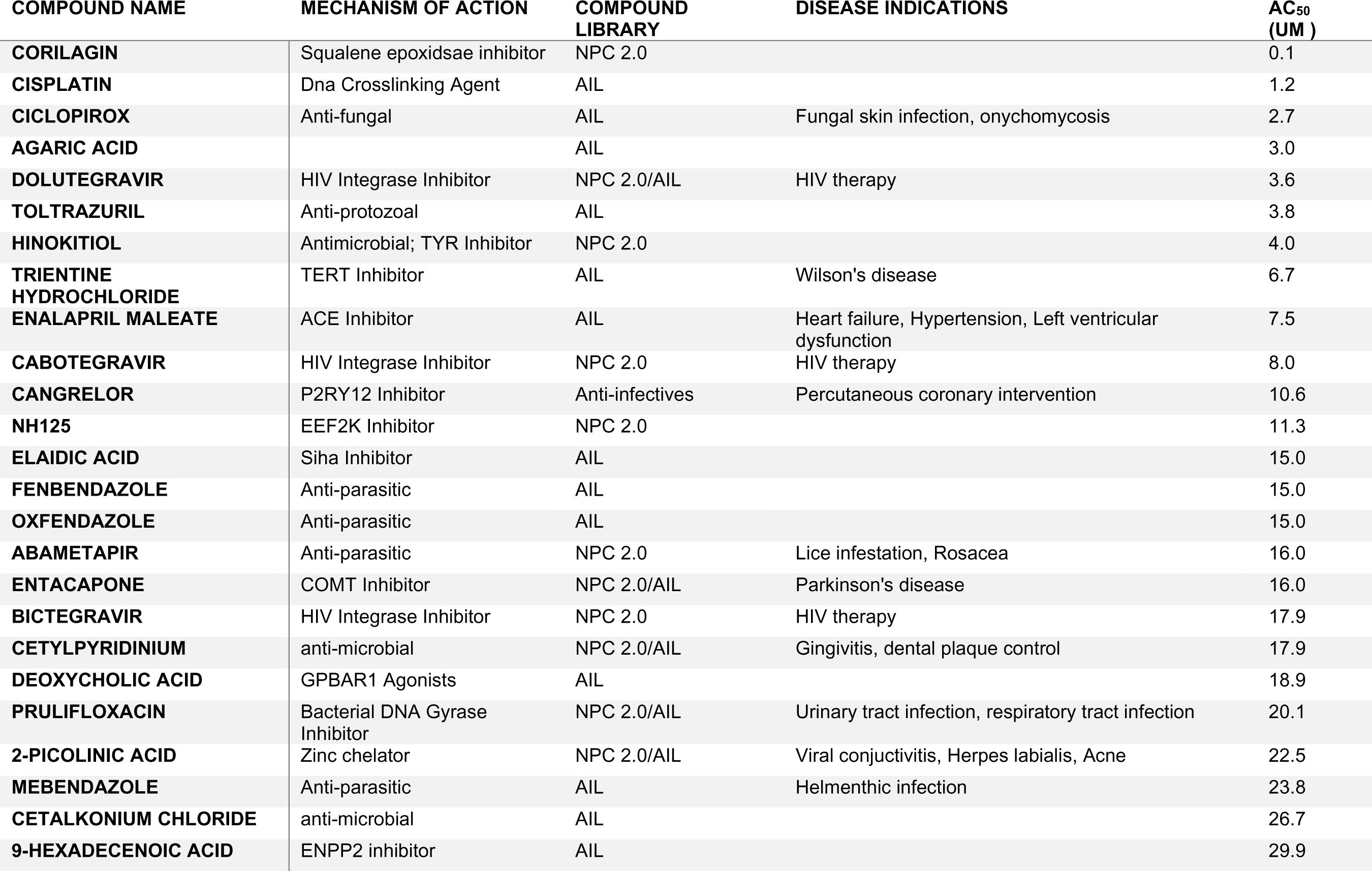
Curated compound actives from the NPC and Anti-Infectives libraries after filtering for assay-interfering compounds. 25 compounds with various MOAs showed weak activity as inhibitors of the ACE2-RBD interaction.

**Figure 4:**
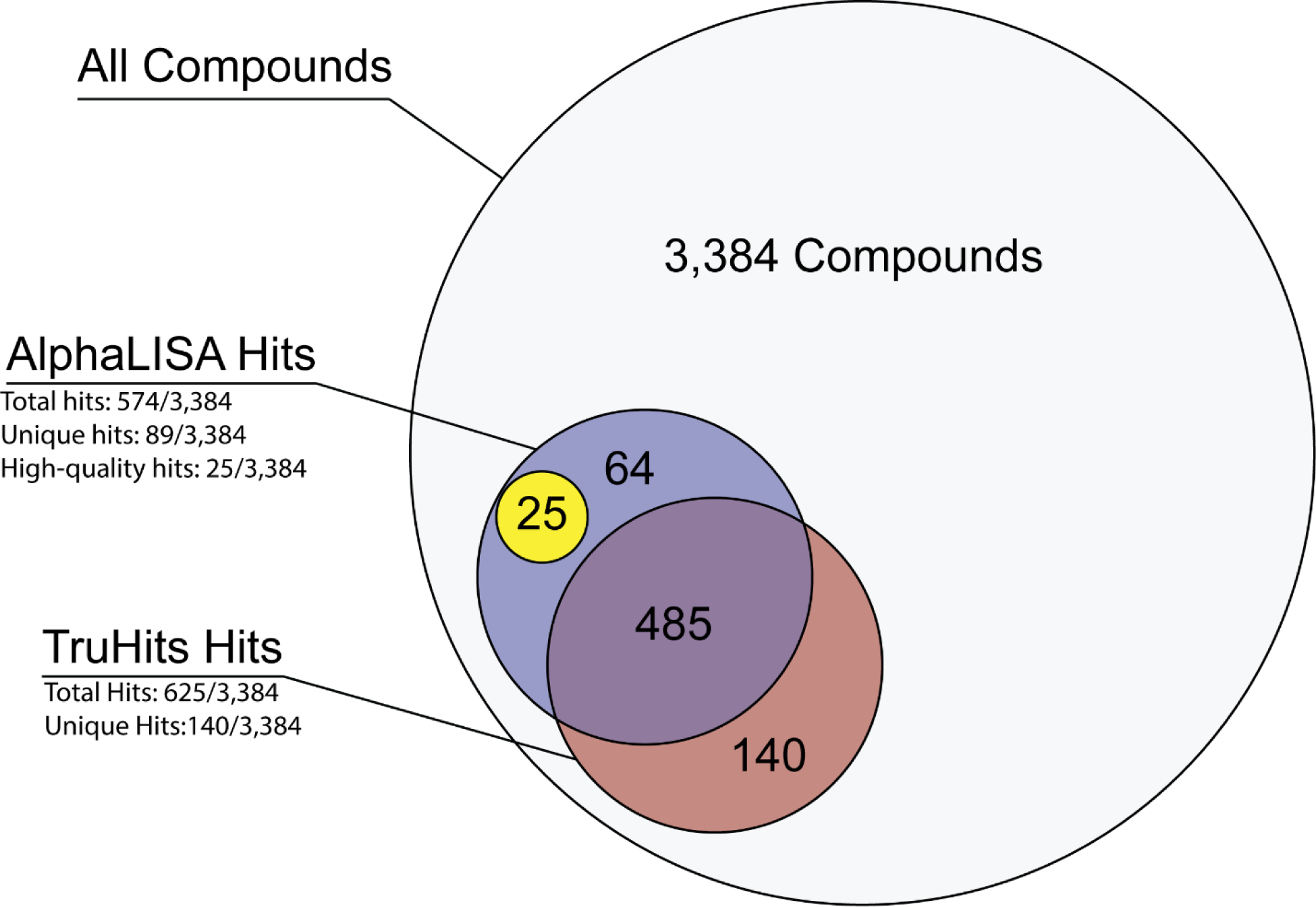
Hit summary from NPC and AIL compound sets. AlphaLISA and TruHits assays were performed against the NPC and AIL compound sets (3,384 compounds total). Potential hits in both the AlphaLISA and TruHits assays were defined as having an IC_50_ ≥ 50 µM and a maximum response ≤ −50%. 89 of those hits were found to be unique to the AlphaLISA assay and those curves were visually inspected to identify 25 high-quality hits. Low-quality hits have TruHits response curves that satisfy the conditions for a “potential hit” but do not significantly diverge from the AlphaLISA data. High-quality hits have minimal TruHits response which do significantly diverge from the AlphaLISA response curve.

**Figure 5:**
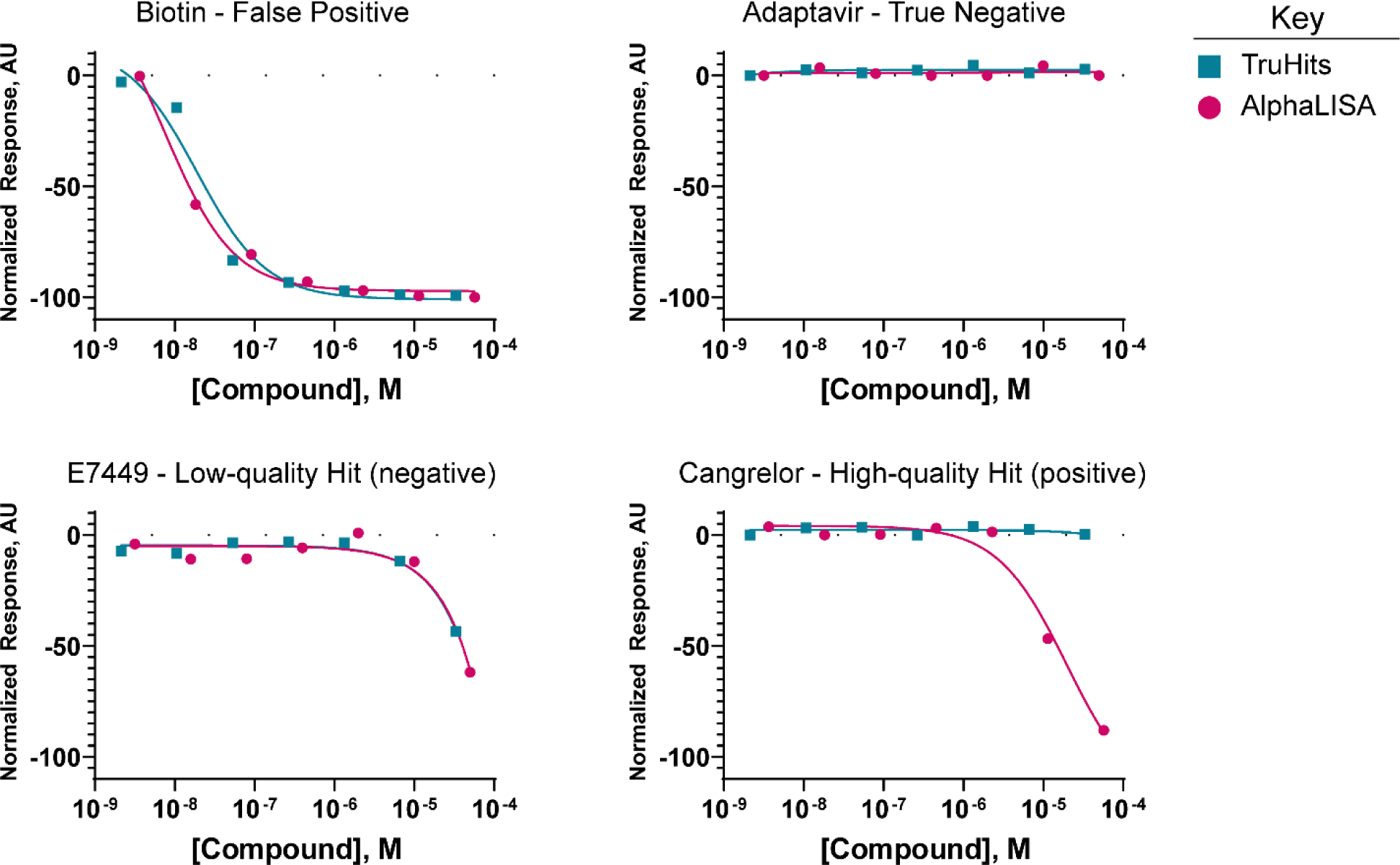
Example dose-response curves showing criteria for hit selections. Biotin is a true false positive active in the AlphaLISA and TruHits assays. Adaptavir, a true negative, was active in neither AlphaLISA nor TruHits Assays. E7449 is a low-quality hit satisfies initial hit identification parameters (IC_50_ ≤ 50 µM and a maximum response ≥50%) but upon visual inspection we see the TruHits data barely miss the initial hit parameters, thus E7449 is not a high-quality hit. Cangrelor is an example of a high-quality hit that satisfies both initial hit identification parameters and passes visual inspection.

## Discussion

We developed the first well-based proximity assay for measuring COVID19 ACE2-RBD interactions using AlphaLISA technology. The AlphaLISA approach has the advantages of high signal-to-background, robust assay performance (Z’ values), and is widely applicable to a variety of therapeutic types (small molecules, nanobodies, peptides, etc.). Molecules which interfere with the generation of or those that scavenge singlet oxygen present a problem as do those which absorb or emit light in the 615-680 nm wavelengths ^20^. However, the ease of use and ability to rapidly deploy an assay using this platform positions it as a strong primary assay for therapeutic screenings. Furthermore, the availability of the TruHits counter-screen simplifies the identification of most false-positive compounds.

While TruHits cannot account for the Protein A-Fc interaction used in our AlphaLISA assay, it does aid in funneling down the potential hits to numbers that are more manageable for lower through-put approaches. Our screen identified 25 high-quality hits that were not active in the counter-assay (Table 3). While there was no expectation of a clinical drug repurposing opportunity emerging from the screen, the hit compounds include several compounds approved for treating patients. A number of molecules containing PAINS moieties such as phenolics appear in Table 1 including the top hit corilagin (a polyphenolic natural product), and careful follow-up would be required to understand mechanism. Nevertheless, the assay can serve to support the identification (HTS) or optimization of small molecule & protein-based therapeutic candidate molecules directed to specifically inhibit the ACE2-spike interaction.

SARS-CoV-2 follows its predecessor, SARS-CoV, by using its spike protein to engage host ACE2 receptors to invade host cells. In response to the SARS-CoV outbreak in 2003 efforts targeting the ACE2-spike interaction emphasized antibody-based therapeutics ^21, 22^. Phage display libraries ^10, 11^ and surface plasmon resonance ^23^ were used to develop neutralizing peptides. However, reported small molecule screens for SARS-CoV at the time focused on target-agnostic cellular approaches ^24^. Focused protein-protein interaction (PPI, proximity) assays were not widely available for screening in 2003. Advances in screening technology over the past decades, such as the AlphaLISA technology used in this assay, enable us to address specific aspects of COVID19 biology on a scale that was inaccessible during previous outbreaks.

Currently, protein-protein interaction assays have been used to determine vital information about ACE2-Spike interactions ^7, 8^, but the methods used (ITC and SPR) are low throughput. At the time of writing, protein-protein interaction assays for ACE2-spike binding have not yet been published, nor had any HTS with ACE2-spike proximity assay approaches. The commercial availability of orthogonal PPI methods using fluorescence resonance energy transfer (FRET), saturation transfer difference (STD) NMR, Co-immunoprecipitation, and microscale thermophoresis support the need for further development of such assays and broad interest of the scientific community.

The AlphaLISA assay we developed presents a quick, simple, and qHTS-amenable approach to ACE2-RBD therapeutic development. However, we are presenting an early assay and orthogonal approaches being developed at NCATS and across the globe can only serve to improve our ability to develop and test COVID19 therapeutics. Indeed, one aspect of ACE2-spike binding we cannot capture in our assay is the role of biological multimerization. Recently multimerization of ACE2 and spike have been demonstrated to enhance the interaction between SARS-CoV-2 spike and host ACE2 ^25^. Future assay development could emphasize the use of additional recombinant protein constructs to more closely model the biological ACES-spike interaction. The advantage to AlphaLISA is that we can continue to use the methods developed herein toward more sophisticated qHTS assay design to suit our evolving knowledge of SARS-CoV-2 biology.

## References

1. Guy, R. K.; DiPaola, R. S.; Romanelli, F.; Dutch, R. E., Rapid repurposing of drugs for COVID-19.Science 2020, 368 (6493), 829–830.

2. Tu, Y. F.; Chien, C. S.; Yarmishyn, A. A.; Lin, Y. Y.; Luo, Y. H.; Lin, Y. T.; Lai, W. Y.; Yang, D. M.; Chou, S. J.; Yang, Y.P.; Wang, M.L.; Chiou, S.H., A Review of SARS-CoV-2 and the Ongoing Clinical Trials. Int J Mol Sci 2020, 21 (7).

3. Eastman, R. T.; Roth, J. S.; Brimacombe, K. R.; Simeonov, A.; Shen, M.; Patnaik, S.; Hall, M. D., Remdesivir: A Review of Its Discovery and Development Leading to Emergency Use Authorization for Treatment of COVID-19. ACS Cent Sci 2020, 6(5), 672–683.

4. Li, G. D.; De Clercq, E., Therapeutic options for the 2019 novel coronavirus (2019-nCoV). Nature Reviews Drug Discovery 2020, 19 (3), 149–150.

5. Lei, C.; Qian, K.; Li, T.; Zhang, S.; Fu, W.; Ding, M.; Hu, S., Neutralization of SARS-CoV-2 spike pseudotyped virus by recombinant ACE2-Ig. Nat Commun 2020, 11 (1), 2070.

6. Li, W. H.; Moore, M. J.; Vasilieva, N.; Sui, J. H.; Wong, S. K.; Berne, M. A.; Somasundaran, M.; Sullivan, J. L.; Luzuriaga, K.; Greenough, T. C.; Choe, H.; Farzan, M., Angiotensin-converting enzyme 2 is a functional receptor for the SARS coronavirus. Nature 2003, 426 (6965), 450–454.

7. Shang, J.; Ye, G.; Shi, K.; Wan, Y. S.; Luo, C. M.; Aihara, H.; Geng, Q. B.; Auerbach, A.; Li, F., Structural basis of receptor recognition by SARS-CoV-2. Nature 2020.

8. Yan, R. H.; Zhang, Y. Y.; Li, Y. N.; Xia, L.; Guo, Y. Y.; Zhou, Q., Structural basis for the recognition of SARS-CoV-2 by full-length human ACE2. Science 2020, 367 (6485), 1444-+.

9. Ho, T. Y.; Wu, S. L.; Chen, J. C.; Wei, Y. C.; Cheng, S. E.; Chang, Y. H.; Liu, H. J.; Hsiang, C. Y., Design and biological activities of novel inhibitory peptides for SARS-CoV spike protein and angiotensin-converting enzyme 2 interaction. Antiviral Res 2006, 69 (2), 70–6.

10. Yeung, K. S.; Meanwell, N. A., Recent developments in the virology and antiviral research of severe acute respiratory syndrome coronavirus. Infect Disord Drug Targets 2007, 7 (1), 29–41.

11. Prabakaran, P.; Zhu, Z.; Xiao, X.; Biragyn, A.; Dimitrov, A. S.; Broder, C. C.; Dimitrov, D. S., Potent human monoclonal antibodies against SARS CoV, Nipah and Hendra viruses. Expert Opin Biol Ther 2009, 9 (3), 355–68.

12. Ju, B.; Zhang, Q.; Ge, J.; Wang, R.; Sun, J.; Ge, X.; Yu, J.; Shan, S.; Zhou, B.; Song, S.; Tang, X.; Yu, J.; Lan, J.; Yuan, J.; Wang, H.; Zhao, J.; Zhang, S.; Wang, Y.; Shi, X.; Liu, L.; Zhao, J.; Wang, X.; Zhang, Z.; Zhang, L., Human neutralizing antibodies elicited by SARS-CoV-2 infection. Nature 2020.

13. Monteil, V.; Kwon, H.; Prado, P.; Hagelkruys, A.; Wimmer, R. A.; Stahl, M.; Leopoldi, A.; Garreta, E.; del Pozo, C. H.; Prosper, F.; Romero, J. P.; Wirnsberger, G.; Zhang, H.; Slutsky, A. S.; Conder, R.; Montserrat, N.; Mirazimi, A.; Penninger, J. M., Inhibition of SARS-CoV-2 Infections in Engineered Human Tissues Using Clinical-Grade Soluble Human ACE2. Cell 2020, 181 (4), 905-+.

14. Brimacombe, K. R.; Zhao, T.; Eastman, R. T.; Hu, X.; Wang, K.; Backus, M.; Baljinnyam, B.; Chen, C. Z.; Chen, L.; Eicher, T.; Ferrer, M.; Fu, Y.; Gorshkov, K.; Guo, H.; Hanson, Q. M.; Itkin, Z.; Kales, S. C.; Klumpp-Thomas, C.; Lee, E. M.; Michael, S.; Mierzwa, T.; Patt, A.; Pradhan, M.; Renn, A.; Shinn, P.; Shrimp, J. H.; Viraktamath, A.; Wilson, K. M.; Xu, M.; Zakharov, A. V.; Zhu, W.; Zheng, W.; Simeonov, A.; Mathé, E. A.; Lo, D. C.; Hall, M. D.; Shen, M., An OpenData portal to share COVID-19 drug repurposing data in real time. bioRxiv 2020, 2020.06.04.135046.

15. Yasgar, A.; Jadhav, A.; Simeonov, A.; Coussens, N. P., AlphaScreen-Based Assays: Ultra-High-Throughput Screening for Small-Molecule Inhibitors of Challenging Enzymes and Protein-Protein Interactions. Methods Mol Biol 2016, 1439, 77–98.

16. Huang, L.; Li, L.; Tien, C.; LaBarbera, D. V.; Chen, C., Targeting HIV-1 Protease Autoprocessing for High-throughput Drug Discovery and Drug Resistance Assessment. Sci Rep 2019, 9 (1), 301.

17. Huang, R.; Southall, N.; Wang, Y.; Yasgar, A.; Shinn, P.; Jadhav, A.; Nguyen, D. T.; Austin, C. P., The NCGC pharmaceutical collection: a comprehensive resource of clinically approved drugs enabling repurposing and chemical genomics. Sci Transl Med 2011, 3 (80), 80ps16.

18. Huang, R.; Zhu, H.; Shinn, P.; Ngan, D.; Ye, L.; Thakur, A.; Grewal, G.; Zhao, T.; Southall, N.; Hall, M. D.; Simeonov, A.; Austin, C. P., The NCATS Pharmaceutical Collection: a 10-year update. Drug Discov Today 2019, 24 (12), 2341–2349.

19. Inglese, J.; Auld, D. S.; Jadhav, A.; Johnson, R. L.; Simeonov, A.; Yasgar, A.; Zheng, W.; Austin, C. P., Quantitative high-throughput screening: A titration-based approach that efficiently identifies biological activities in large chemical libraries. P Natl Acad Sci USA 2006, 103 (31), 11473–11478.

20. Coussens, N. P.; Auld, D.; Roby, P.; Walsh, J.; Baell, J. B.; Kales, S.; Hadian, K.; Dahlin, J. L., Compound-Mediated Assay Interferences in Homogenous Proximity Assays. In Assay Guidance Manual, Sittampalam, G. S.; Grossman, A.; Brimacombe, K.; Arkin, M.; Auld, D.; Austin, C. P.; Baell, J.; Bejcek, B.; Caaveiro, J. M. M.; Chung, T. D. Y.; Coussens, N. P.; Dahlin, J. L.; Devanaryan, V.; Foley, T. L.; Glicksman, M.; Hall, M. D.; Haas, J. V.; Hoare, S. R. J.; Inglese, J.; Iversen, P. W.; Kahl, S. D.; Kales, S. C.; Kirshner, S.; Lal-Nag, M.; Li, Z.; McGee, J.; McManus, O.; Riss, T.; Saradjian, P.; Trask, O. J., Jr.; Weidner, J. R.; Wildey, M. J.; Xia, M.; Xu, X., Eds. Bethesda (MD), 2004.

21. Sui, J. H.; Li, W. H.; Murakami, A.; Tamin, A.; Matthews, L. J.; Wong, S. K.; Moore, M. J.; Tallarico, A. S. C.; Olurinde, M.; Choe, H.; Anderson, L. J.; Bellini, W. J.; Farzan, M.; Marasco, W. A., Potent neutralization of severe acute respiratory syndrome (SARS) coronavirus by a human mAb to S1 protein that blocks receptor association. P Natl Acad Sci USA 2004, 101 (8), 2536–2541.

22. Zhu, M., SARS Immunity and Vaccination. Cell Mol Immunol 2004, 1 (3), 193–8.

23. Struck, A. W.; Axmann, M.; Pfefferle, S.; Drosten, C.; Meyer, B., A hexapeptide of the receptor-binding domain of SARS corona virus spike protein blocks viral entry into host cells via the human receptor ACE2. Antivir Res 2012, 94 (3), 288–296.

24. Kao, R. Y.; Tsui, W. H.; Lee, T. S.; Tanner, J. A.; Watt, R. M.; Huang, J. D.; Hu, L.; Chen, G.; Chen, Z.; Zhang, L.; He, T.; Chan, K. H.; Tse, H.; To, A. P.; Ng, L. W.; Wong, B. C.; Tsoi, H. W.; Yang, D.; Ho, D. D.; Yuen, K. Y., Identification of novel small-molecule inhibitors of severe acute respiratory syndrome-associated coronavirus by chemical genetics. Chem Biol 2004, 11 (9), 1293–9.

25. Lui, I.; Zhou, X. X.; Lim, S. A.; Elledge, S. K.; Solomon, P.; Rettko, N. J.; Zha, B. S.; Kirkemo, L. L.; Gramespacher, J. A.; Liu, J.; Muecksch, F.; Lorenzi, J. C. C.; Schmidt, F.; Weisblum, Y.; Robbiani, D. F.; Nussenzweig, M. C.; Hatziioannou, T.; Bieniasz, P. D.; Rosenburg, O. S.; Leung, K. K.; Wells, J. A., Trimeric SARS-CoV-2 Spike interacts with dimeric ACE2 with limited intra-Spike avidity. bioRxiv 2020, 2020.05.21.109157.

